# ViraMiner: Deep Learning on Raw DNA Sequences for Identifying Viral Genomes in Human Samples

**DOI:** 10.1101/602656

**Authors:** Ardi Tampuu, Zurab Bzhalava, Joakim Dillner, Raul Vicente

## Abstract

Despite its clinical importance, detection of highly divergent or yet unknown viruses is a major challenge. When human samples are sequenced, conventional alignments classify many assembled contigs as “unknown” since many of the sequences are not similar to known genomes. In this work, we developed ViraMiner, a deep learning-based method to identify viruses in various human biospecimens. ViraMiner contains two branches of Convolutional Neural Networks designed to detect both patterns and pattern-frequencies on raw metagenomics contigs. The training dataset included sequences obtained from 19 metagenomic experiments which were analyzed and labeled by BLAST. The model achieves significantly improved accuracy compared to other machine learning methods for viral genome classification. Using 300 bp contigs ViraMiner achieves 0.923 area under the ROC curve. To our knowledge, this is the first machine learning methodology that can detect the presence of viral sequences among raw metagenomic contigs from diverse human samples. We suggest that the proposed model captures different types of information of genome composition, and can be used as a recommendation system to further investigate sequences labeled as “unknown” by conventional alignment methods. Exploring these highly-divergent viruses, in turn, can enhance our knowledge of infectious causes of diseases.

## 1 Introduction

The human virome is the collection of all viruses that reside on a human body. Many different viruses are present in human samples and their composition appears to be different in diseased individuals[1, 2]. Despite its clinical importance, its full impact on human health is not fully understood [3, 4] and the detection and classification of human viruses represents a major challenge.

Current metagenomic studies detect many novel viruses, which indicates that only a small part of human viruses has been discovered and many others are yet to be reported[5, 6, 7, 8, 9, 10]. Studies report epidemiological indications that there may exist undiscovered pathogens. For example, there is correlative evidence suggesting that the viruses may be involved in the development of autoimmune diseases such as diabetes[11] and multiple sclerosis[12].

Next Generation Sequencing (NGS) technologies provide a powerful tool to directly obtain the DNA sequences present in clinical samples without any prior knowledge of the biospecimens[13]. The term metagenomics implies complete sequencing and recovering of all non-human genetic material that might be present in a human sample and accordingly, analytical techniques for virus discovery are commonly applied to metagenomic samples[5, 7, 8, 14, 15, 16, 17, 18, 19].

Currently, the detection of potential viral genomes in human biospecimens is usually performed by NCBI BLAST, which implements alignment-based classification where sequences are aligned to known genomes from public databases and then estimates how much percentage similarity they share. However, metagenomic samples might contain a large number of highly divergent viruses that have no homologs at all among known genomes. As a consequence, many sequences generated from NGS technologies are classified as “unknown” by BLAST[5, 18]. Another commonly used algorithm for viral discovery in metagenomic samples is HMMER3[20], which applies profile Hidden Markov Models (pHMM) in relation to the vFams[21] database of viral family of proteins, that have been built by multiple sequence alignments from all viral proteins in RefSeq. With this method, sequences are compared to entire viral families which enables it to be more effective in detecting distant homologs[22]. However, this pipeline needs a reference database which is a drawback while conducting the analysis of identification of highly divergent viruses. A different approach consists of using machine learning techniques to learn from examples to classify viral genomes and to generalize to novel samples. In particular, several machine learning models for detecting viruses in metagenomic data have been already published[23, 24, 25], but none of them were trained nor tested to identify viruses in different human biospecimens.

In this study, we have developed ViraMiner, a methodology which uses Convolutional Neural Networks (CNN) on raw metagenomic contigs to identify potential viral sequences across different human samples. The architecture of ViraMiner builds on top of the CNN architecture of [25], adding major extensions for it to be more effective on this particular classification problem. For training the model we used 19 metagenomic experiments originating from sample types such as skin, serum, and condylomata. The model achieves a significantly improved accuracy compared to other existing methods for virus identification in metagenomic samples. To our knowledge, the proposed model is the first methodology that can detect the presence of viruses on raw metagenomic contigs from various human biospecimens. A pre-trained model ready to be used is also publicly available on GitHub (https://github.com/NeuroCSUT/ViraMiner).

## 2 Results

First we will describe the overall architecture of the proposed model (ViraMiner) for the detection of viral genomes from assembled metagenomic contigs (see Methods for a detailed description of the architecture, its inputs and outputs and detail on training procedure). In the following sections, performance measures of the model and multiple baselines are reported. Finally, an evaluation of the model generalization when applied to novel metagenomic experiments is given.

### 2.1 *ViraMiner* architecture

In this work we developed ViraMiner, a CNN-based method for detecting viral contigs in human metagenomic datasets. As illustrated on Figure 1, ViraMiner receives a raw DNA sequence as input and outputs a single value that indicates the likelihood of this sequence being of viral origin. In ViraMiner we apply convolutional layers with learnable filters straight on the raw sequences. This way the model has access to all information and can learn by itself from labeled examples which features are important to extract (which information to keep).

**Figure 1:**
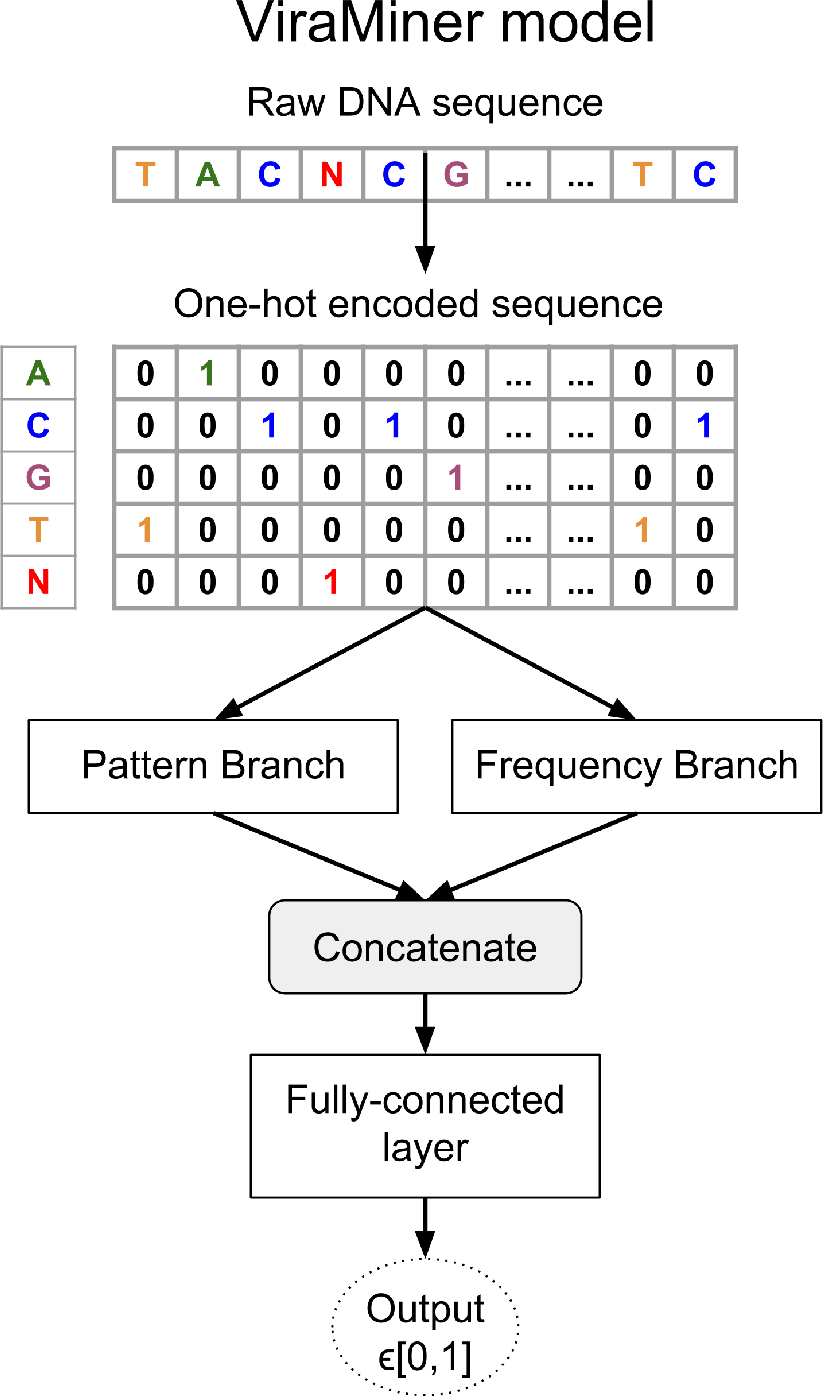
ViraMiner architecture. ViraMiner architecture takes as input the raw sequences in one-hot encoded form. The raw sequences are then processed by two different convolutional branches. The outputs of the branches are joined and used for computing the final output. The output value is restricted to range [0,1] and reflects the likelihood of the sequence belonging to virus class.

To maximize the model’s ability to extract relevant features, the ViraMiner model has two different convolutional branches (see Figure 1). Intuitively, one of the branches (pattern branch) returns how well certain patterns were maximally matched along the DNA sequence. In this branch the convolutional layer is followed by *global max pooling* that outputs only the maximal value for each convolutional filter. The other (frequency branch) returns information about pattern frequencies. In this branch the convolutional layer is followed by *gloabl average pooling* that yields the average value for each convolutional filter. The final result is linearly decoded from the joint output of these two branches.

### 2.2 Identifying viral genomes in human samples

The proposed models were trained on metagenomic assembled contigs from 19 different experiments, which in turn were sequenced from different human sample types (serum, skin, condylomata, …). These contigs were combined, shuffled, and partitioned into training (80%), validation (10%), and testing (10%) sets. Models were trained using the training set and the best model was selected according to validation set performance (see “Hyperparameter search and testing” subsection in Methods). The test set was used only for the final evaluation of the best models.

The proposed architecture achieved outstanding test performance on this human metagenomic data. On Fig 2 we show the Receiver Operating Characteristic (ROC) curve of the best ViraMiner model with an area under the curve of 0.923.

**Figure 2:**
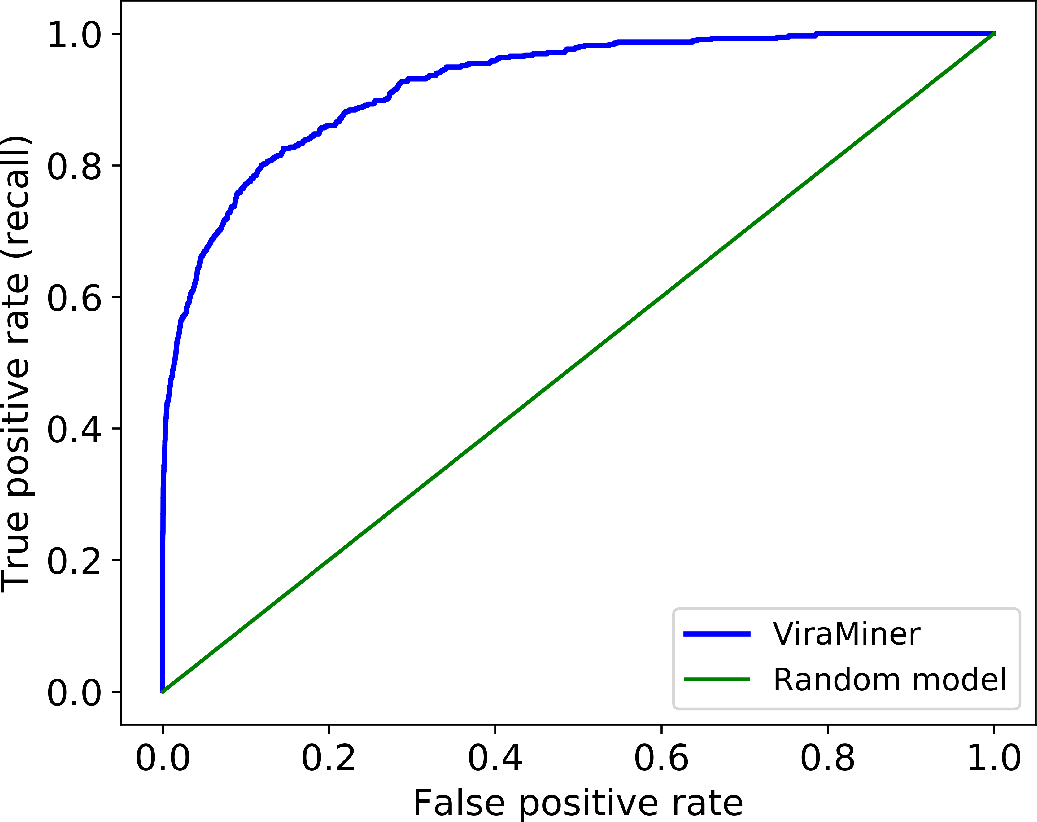
ROC curve of ViraMiner model on the testing dataset. The blue line on the figure represents the performance of ViraMiner (AUROC: 0.923). For comparison, the green line depicts the performance of a random model.

Given the huge class imbalance of these type of datasets (e.g. in the studied human metagenomics dataset there were only 2% of viral contigs) it is important to evaluate precision and recall measures for both classes separately. Overall measures are hardly informative - a trivial model classifying everything as non-virus would achieve 98% overall precision and 98% overall recall, but it would nevertheless be useless for detecting viral genomes. Thus, next we report the precision and recall of the virus class instead of global performance. Note also that to measure particular recall and precision values, one would need to establish a classification threshold. The higher the threshold is set, the higher the precision gets but at the expense of recall. On the other hand, setting a lower threshold would increase recall but decrease precision. Fig 3 represents the tradeoff between precision and recall for the virus class. The dotted lines indicate that ViraMiner could achieve 90% precision with 32% recall. With a stricter threshold, one can obtain 95% precision and 24% recall. Inversely, if one wishes to increase recall and accept relatively more false positives, the model can reach 41% recall with 70% precision.

**Figure 3:**
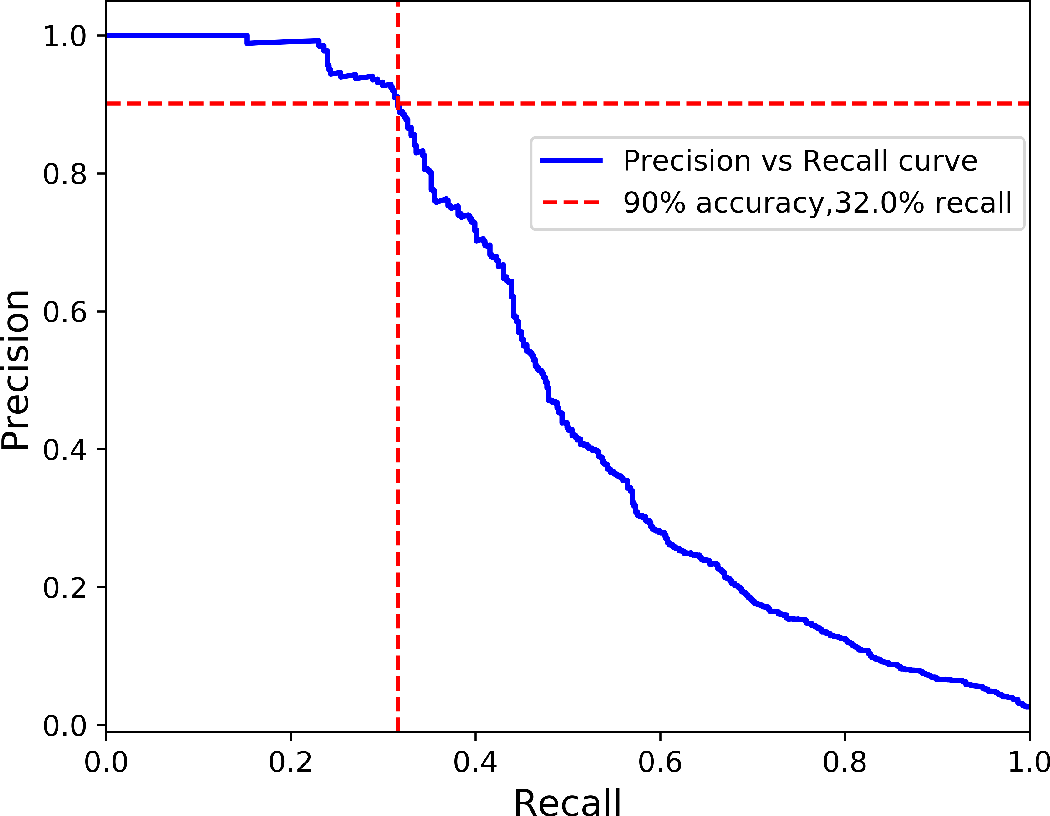
The trade-off between precision and recall for classifying viral genomes. A genome is labeled as virus if the output value is higher than a certain threshold. The blue line represents the trade-off. The crossing point of the vertical and horizontal red dotted lines indicates that the model can achieve 90% precision with 32% recall.

### 2.3 Baselines comparison

#### 2.3.1 Contributions of each CNN branch separately

The model training proceeds by training the Pattern and Frequency branches separately. Consequently, we can also report the performance of each branch alone. With hyperparameters selected to maximize AUROC on validation set, on test set the Pattern branch achieves AUROC 0.905. In the same conditions, the Frequency branch achieves a test AUROC of 0.917. This shows that the filters of each branch result in a relatively high accuracy despite their different pooling operations, and their combination results in an even higher AUROC score (0.923). The stepwise training of ViraMiner architecture is important to avoid overfitting. When optimizing altogether the 2 million parameters in a ViraMiner model at the same time (both branches trained simultaneously), the performance on test set decreases. The test AUROC of such end-to-end trained model is only 0.896.

#### 2.3.2 K-mer models

Most of the previous machine learning models for classifying metagenomic datasets are based on k-mer counting [24, 26, 27]. To compare against the performance of such methods, we also extracted k-mers from the investigated dataset and trained Random Forest (RF) classifiers on the extracted values, while keeping the same data partitioning as above. RF is a competitive machine learning algorithm for non-linearly separable classes and it has already been used on this type of datasets [23, 28]. The best test performance with RF models was achieved with 6-mers and it produced test AUROC 0.875 (Fig 4). RF performances on 3- 4- 5- and 7-mers were 0.867, 0.872, 0.873, and 0.869 respectively (Fig 5).

**Figure 4:**
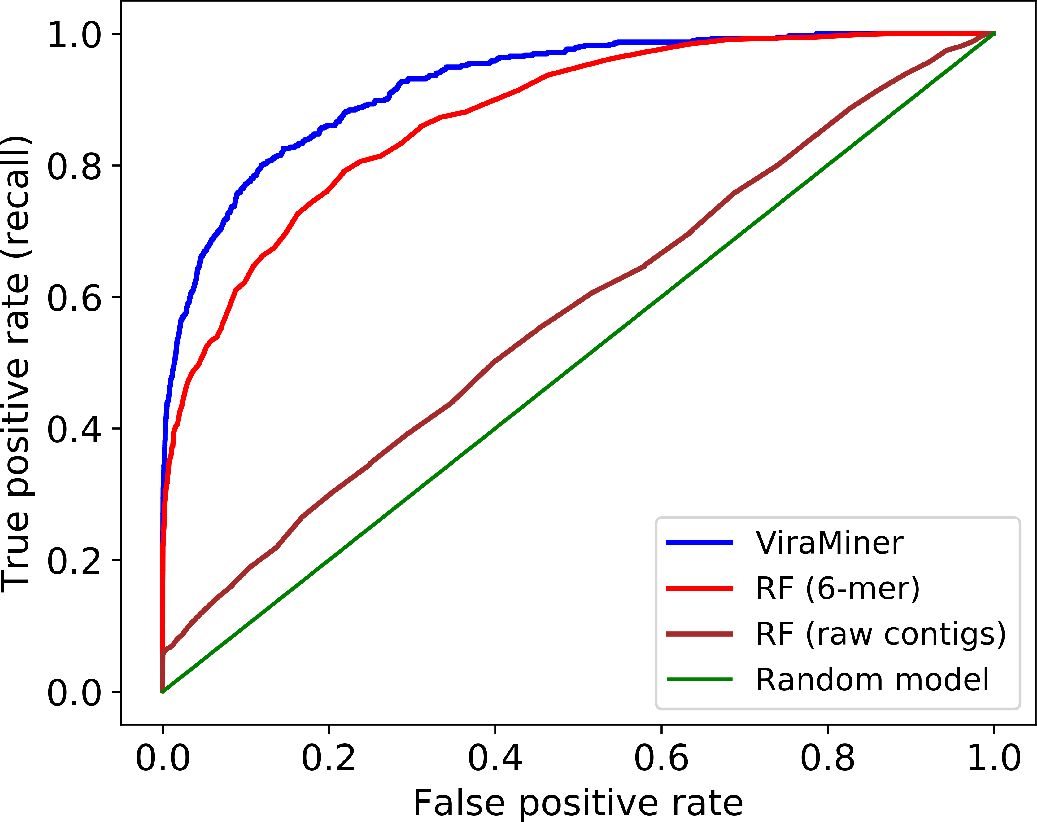
Comparison with baseline models. The blue line depicts the performance of ViraMiner (AUROC: 0.923); the red line corresponds to the best of Random Forest on k-mers models (using 6-mers); the brown line, shows RF performance on raw metagenomic contigs.

**Figure 5:**
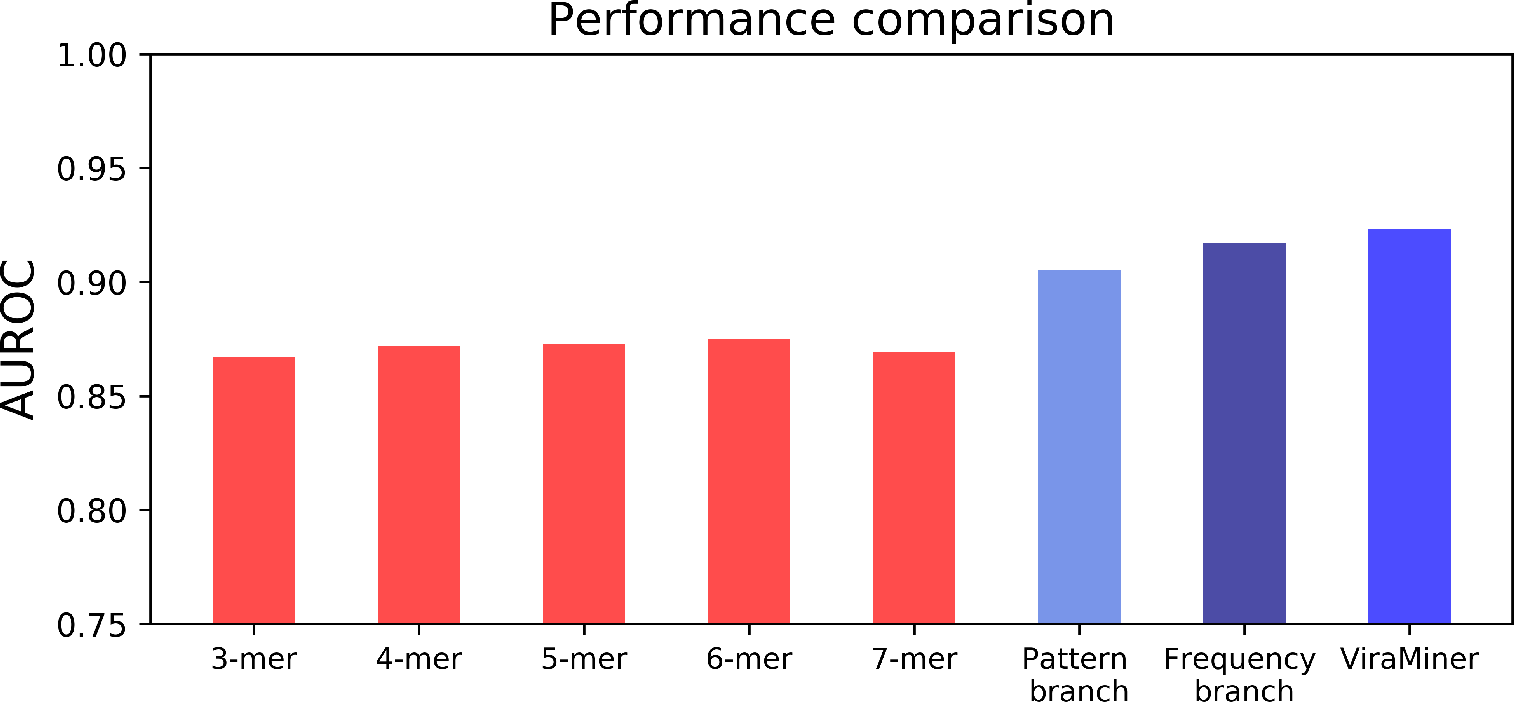
Performance comparison of models designed in this study. Models based on k-mers (red bars) were trained using Random Forest and the best performance was achieved with 6-mers with test AUROC value 0.875. Convolutional Neural Networks (blue bars) use the raw sequence as input and outperform RF models. Pattern and Frequency branches yield performances above 0.9 AUROC. ViraMiner uses a combination of both branches and achieves the highest test AUROC value (0.923) out of all models.

For completeness, we also trained a RF model on raw metagenomic contigs (one-hot encoded and flattened), that is the same input used to ViraMiner, and hence without the manual extraction of different k-mers counts. The test performance of such RF model was very close to a random model (AUROC: 0.573).

The overall comparison of test performances of models designed in this study is summarized in Fig 5.

#### 2.3.3 Positive Predictive Values

We also compared Positive Predictive Value (PPV) of ViraMiner with PPV of the best RF (6-mer) model. Top 20 test set sequences most likely to be viruses according to both models were indeed viruses. However, there was a clear difference between confidence rates: confidence of ViraMiner for its top 20th virus was 99% while RF yielded only 67% for its corresponding virus on the 20th position. For the top 200 sequences, ViraMiner achieved 88% PPV whereas RF reached 75%. Pattern and Frequency branches for the same number of top sequences output 79% and 83% PPVs respectively.

### 2.4 Performance on previously unseen sequencing experiments

In this section we describe the performance of the proposed model on novel metagenomic experiments. We retrain the model with the same hyperparameters as above (no new search is performed), but we remove one sequencing experiment completely from the train and validation sets and use it as the test set. Notice that the training procedure remains absolutely the same - we train the models (branches first, then the full model) until validation AUROC stops increasing. Afterwards, the model is applied to the test set - the left-out data originating from a novel sequencing experiment. To show dataset-to-dataset variability, we repeat this leave-one-experiment-out procedure for all metagenomic datasets originating from serum sample type (see Table 1). By choosing these experiments for the leave-out procedure we also check if the model can generalize on one specific sample type. To run these tests we selected serum because in our dataset largest number of experiments and largest number of samples originate from this sample type.

**Table 1:** ViraMiner Performance on unseen serum metagenomic experiments. The number of viral samples in these experiments varies from 22 to 732.

| Left-out experiment | 2011_G5 | 2014_G1 | 2014_G5 | 2014_G6 | 2014_G7 | Micro-average |
| --- | --- | --- | --- | --- | --- | --- |
| test AUROC | 0.95 | 0.89 | 0.96 | 0.92 | 0.86 | 0.94 |

The ViraMiner model achieved good test performance on all 5 serum datasets, with AUROCs ranging from 0.86 to 0.96. To provide a unique measure, we merge the predictions for all five sets and redraw the ROC curve. The area under this micro-averaged ROC curve across the five test sets is 0.94.

We also investigated whether training a ViraMiner model on data originating from only serum metagenomic datasets would improve the performance compared to a model trained using a variety of metagenomic data sets (as above). For this purpose, we trained ViraMiner model on assembled contigs originating only from four serum experiments and tested on the 5th serum dataset. We repeated to the leave-out procedure five times, so each dataset was left out once. With these reduced but more specific training set, the test area under micro-averaged ROC curve was 0.91. This is a noticeable decrease compared to 0.94 AUROC achieved when using all available training data. In our case using all available data points was useful even though many of these data points came from completely different sample types such as skin, cervix tissue, prostate secretion, etc.

## 3 Discussion

In this work, we proposed ViraMiner, a CNN-based method to detect viruses in diverse human biospecimens. The tool includes two different branches of CNN: a Pattern branch with convolution+max operator and a Frequency branch with convolution+average operator.

Firstly, we trained the model on assembled metagenomic contigs originating from 19 metagenomic experiments which were merged and partitioned into training, validation and testing sets. The model achieves 0.923 test area under the ROC curve. This is a significant improvement compared to a previous tool ([28], reaching 0.79 AUROC) that was designed using relative synonymous codon usage (RSCU) and trained on data originating from the same metagenomic experiments. We are confident to conclude that using raw metagenomic contigs with CNN can extract much more information for the problem of viral classification. Furthermore, another advantage of ViraMiner is that a sequence is not required to have an Open Reading Frame (ORF) which is a central requirement for the models based on codon usage.

VirFinder and DeepVirFinder (DVF) are other similar tools available for detecting viruses in metagenomic sequencing datasets [24, 25]. Former is based on k-mers while latter applies CNN on raw DNA sequences. These tools, however, were specifically trained to identify prokaryotic viruses whereas we trained the model to detect any virus that might appear in multiple human samples. The architecture of ViraMiner builds on top of the CNN architecture of DeepVirFinder, adding major extensions for it to be more effective on this particular classification problem. Indeed, the newly introduced Frequency model separately performed better than the DVF architecture (referred to as Pattern model in Results). Considering these two studies, DVF and the present article, it is clear that such CNN architectures work very well on viral genome classification.

We also explored whether the proposed CNN architecture was able to extract more informative features from raw metagenomic contigs than to just count k-mers on them. We counted 3-,4-,5-,6- and 7-mers on the contigs, trained Random Forest and compared the results with our model. Since ViraMiner produced a higher accuracy (Fig 4) we can deduce that the architecture extracts more complex features of genome compositions that can detect patterns that cannot be identified with just counting k-mers.

The most important criteria that ViraMiner had to satisfy, however, was to generalize its classification abilities on completely new and unseen metagenomic experiments from which the model had not seen any data point. To this end, not only did we test ViraMiner on one specific experiment but also one specific sample type. In our dataset, the largest number (five) of experiments came from serum and therefore, we tested if the model could generalize on all these five experiments. ViraMiner was retrained five times with the same hyperparameters but each time data from one experiment was left out and used as the test set. The model produced slightly different test AUROC values for each of these datasets which gave in an average 0.94 area under ROC curve.

Certainly, at this stage ViraMiner or other deep learning based methods cannot replace BLAST or HMMER3 because of the limited number of data points in the training dataset. However, it is important to note that ViraMiner captures different kind of information of genome composition. Considering the results described above, the proposed method can be used as a recommendation system to further research “unknown” sequences labeled by the conventional methods among which highly divergent viruses may be hidden. Pre-trained models or ViraMiner are publicly available at *https://github.com/NeuroCSUT/ViraMiner*. We hope that this and future methods can discover biological attributes of viral genome to predict and reveal viruses, and in turn, help focus our exploration of infectious causes in diseases.

## 4 Methods and Materials

### 4.1 Samples and sequencing types

The metagenomic sequences in this work were generated using Next Generation Sequencing platforms such as the NextSeq, Miseq, and HiSeq (Illumina) as described by the manufacturer. The dataset was derived from human samples belonging to different patient groups that have been described in detail in[6, 7, 8, 29, 30, 31]. The goal of those analyses was to detect viral genomes or other microorganisms in diseased individuals or in their matched control subjects.

### 4.2 Bioinformatics

All the sequencing experiments were processed and analyzed using a benchmarked bioinformatics workflow[32]. To summarize, we start analysis with quality checking where reads are filtered based on their Phred quality score. Afterwards, quality checked reads that align to human, bacterial, phage and vector DNA with >95% identity over 75% of their length are subtracted from further analysis using BWA-MEM[33]. The reads that are left are normalized and then assembled using the IDBA-UD[34], Trinity[35], SOAPdenovo, and SOAPdenovo-Trans[36] assemblers. The assembled contigs are then subjected to taxonomic classification using BLAST. The code of the pipeline is available on GitHub (https://github.com/NGSeq/ViraPipe and https://github.com/NIASC/VirusMeta).

### 4.3 Data processing and labeling

The training dataset included 19 different NGS experiments that were analyzed and labeled by PCJ-BLAST [37] after applying the de novo genome assembly algorithms. Parameters for PCJ-BLAST were as follows: type of algorithm=Blastn; nucleotide match reward =1; nucleotide mismatch penalty =1; cost to open a gap =0; cost to extend a gap =2; e-value ≤ *e*^−4^. All assembled contigs that were labeled by this bioinformatics workflow were merged to train several machine learning algorithms.

To extract input dataset for Convolutional Neural Networks, labeled sequences were divided into equal pieces (300bp and 500bp), each one of them was labeled as the original sequence and remaining nucleotides at the end of the contigs were removed from the further analysis. For example, according to this processing 650 bp viral contig would produce two equal 300bp viral sequences and the last 50 nucleotides would be removed from the input dataset. Initially, we generated two training datasets based on sequence lengths: first with 300bp and the second with 500bp. After noticing that 300bp contigs producing significantly better results we continued working only on this dataset and dropped the other. Furthermore, all contigs that contained even one “N” letter (reference to any nucleotide) were removed from the further analysis.

For computing baselines, we also counted k-mers in the processed dataset. Given that extracting k-mers becomes more and more computationally expensive as k increases, we conducted distributed and parallel computing of k-mers by using Hadoop (https://hadoop.apache.org) and Spark[38]. The Spark code (available under https://github.com/NIASC/ViraPipeV2) outputs 4^*k*^ × *l* matrix where *l* is number of rows from an input dataset.

### 4.4 Machine Learning Methods

#### 4.4.1 Convolutional Neural Networks (CNNs)

Convolutional neural networks(CNNs) are a type of feedforward neural networks that learn from example input-output pairs via gradient descent[39, 40]. CNNs are most widely used in image processing[39, 41], but have been successfully applied in various fields[42, 43], including analysis of biological sequences such as DNA and amino acid sequences[44, 45, 46]. The convolutional layers treat their inputs as multidimensional arrays - an image is treated as a 2D array with 3 color channels, while a DNA sequence is seen as a 1D array with one channel per nucleotide. In the present work we use sequences of length 300 and consider 5 possible values (ATGCN) at each position, corresponding to a 1D sequence of length 300 with 5 channels. A convolutional layer is characterized by a set of learnable filters that are convolved along the input DNA sequence (or along the width and height of an image). At each position the filter is applied to, a dot product between the weights of the filter and the input values is computed. The dot products resulting from applying the same filter to different positions along the sequence form a feature map. A convolutional layer usually has many filters and hence its output is a set of feature maps. On this output, further computations can be performed such as applying an activation function, pooling or adding further layers. The parameter values in the filters, like all other learnable parameters in the model, are optimized by the gradient descent algorithm.

#### 4.4.2 CNN architectures for predicting the viral nature of sequences

In this work we use a convolutional neural network called ViraMiner to predict whether a given DNA sequence is of a viral origin or not. This model contains two convolutional branches that contribute different types of information to the final fully-connected decision-making layer (Fig6a).

**Figure 6:**
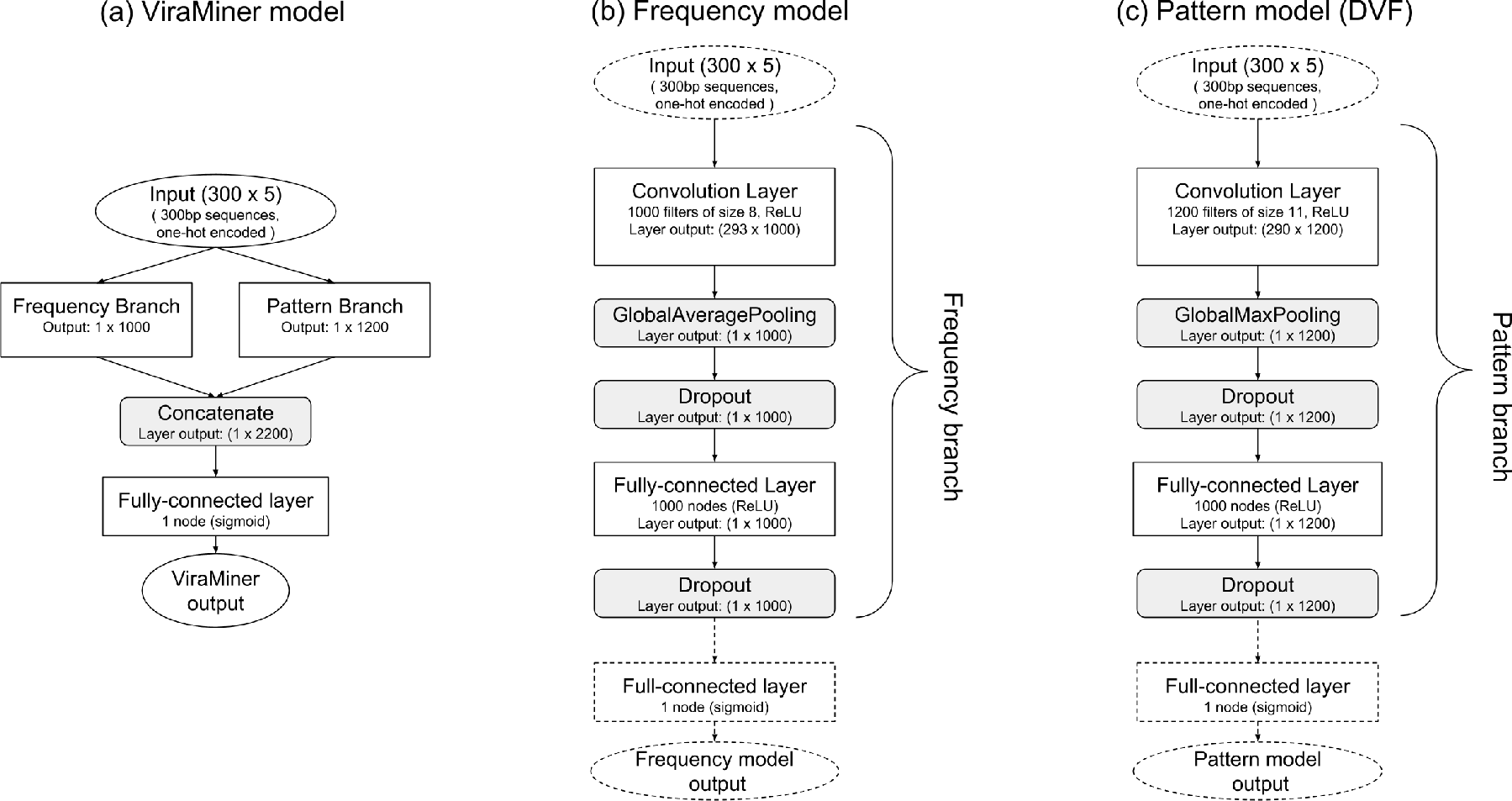
ViraMiner architecture. (a) ViraMiner full model, with the branches not expanded. (b) Architecture of the Frequency model with best performing layer sizes shown. The dotted parts will be discarded when using the pre-trained parameter values of this model as the Frequency Branch in the full model. (c) Architecture of the Pattern model (similar to DeepVirFinder) with best performing layer sizes shown. The dotted parts will be discarded when using the pre-trained parameter values of this model as the Pattern Branch in the full model.

This model is partly based on the DeepVirFinder (DVF) architecture by Ren et al[24]. The DVF model is a Convolutional Neural Network (CNN) that applies the *max operator* on the outputs of each convolutional filter (on each feature map). Only the one maximal activation value per filter is passed on to the next layers. All other information, such as how often a good match was found, is lost. This architecture is illustrated on Fig6c.

In this work we suggest that a convolutional layer followed by *average operator* instead of *max operator* provides important information about the frequency of patterns, information that is otherwise lost. In such a case, while losing information about the maximal activation (best match), we gain information about frequency-the average cannot be high if only a few good matches were found. In previous work the authors of DVF and the authors of the current article have shown that methods based on pattern frequency (k-mer counts, relative synonymous codon usage) are effective in separating viral samples from non-viral ones [24, 28]. Using *convolution + average* is a natural extension to these pattern counting-based models. This results in the architecture on Fig6b.

To clarify, we do not claim that using *convolution+max* is without merit, it simply captures a different type of information. In fact, in our ViraMiner model we combine feature maps processed by averaging and by max operators. This allows the model to base its decisions on both pattern-matching and pattern-frequency information. The ViraMiner detailed architecture is shown on Fig6a where Frequency Branch and Pattern Branch refer to the architectures on Fig6b and Fig6c, without the input and output layers.

#### 4.4.3 Training

It is important to note that Pattern and Frequency models can be used as a separate models and trained independently. In our training scheme we first train the two models, then remove the output layers and use the middle layers as Frequency and Pattern branches in the full ViraMiner model. We finally optimize only the parameters in the final layer of the ViraMiner architecture, leaving the weight and bias values in the branches unchanged, exactly as they were in the independent models. Notice that in this last step of learning, values of only roughly two thousand parameters are changed.

Compared to training the ViraMiner architecture end-to-end, that is without pre-training and freezing the branches, this step-wise training procedure helps avoid overfitting. A ViraMiner model where all parameters are trained together from scratch overfits and performs significantly worse on unseen data. Similarly, overfitting increases when initializing the branches with pre-trained values, but fine-tuning them when training the full ViraMiner model.

#### 4.4.4 Hyperparameter search and testing

We merged and shuffled the data originating from 19 metagenomics sequencing experiments and divided it into training (80%), validation (10%) and test sets (10%). We then performed an extensive hyperparameter search for the Pattern and Frequency models separately. For both architectures, hundreds of variants were trained using the 80% of data in training set. The performance of these models was measured by AUROC on validation set. The models were trained until validation AUROC had not increased for 6 consecutive epochs, but not longer than 30 epochs. After each epoch the model was saved only if validation AUROC had increased. Adam optimizer [47] and batch size 128 were used in all cases.

The initial hyperparameter search scanned the following parameters:

- Filter size: in range from 4 to 40 with step of 4.
- Learning rate: 0.01, 0.001 and 0.0001. Each value was tried with and without learning rate decay, which multiplies learning rate by 0.5 after each 5 epochs.
- Layer sizes (applies both to number of filters in convolutional layers and to number of nodes in fully connected layer): 1000, 1200 or 1500
- Dropout probability: 0.1 or 0.5

In a second stage of hyperparameter search, each possible filter size value (with step of 1) was scanned in the region of interest determined by initial scan (8 to 16 for Frequency model; 6 to 14 for Pattern model). For other hyperparameters, some values were left out from the second stage as they clearly worsened the performance.

When the extensive hyperparameter scan was completed, the best Pattern model and best Frequency model (as per validation AUROC) were selected and used to initialize the branches in the ViraMiner architecture. We then trained the ViraMiner model with different learning rates, learning rate decays and with/without fine-tuning the branches and again selected the best model according to validation AUROC.

Having selected best models according to validation performance, we finally evaluated the best Pattern, best Frequency and best ViraMiner model on the test set which had been left unused up until this point.

#### 4.4.5 Baselines

Using 80/10/10 split, we also trained two types of baseline models. First of all, using k-mer counts extracted from the contigs as input, we trained Random Forest models with 1000 trees (all other parameters as default in scikit-learn Python package[48]) for k in range from 3 to 7. In results section, we report each model’s test AUROC.

Secondly, we also trained a Random Forest model directly on the 300bp sequences, without any feature extraction. For that we one-hot encoded the sequences and flattened them, resulting in 1200 inputs, 4 per each position in the sequence. Notice that such model is not position invariant - a shift of the sequence by just one base pair would completely change the output.

### 4.5 Code availability

All code, for ViraMiner model and for baselines, was written using Python2.7. For neural networks, Keras library was used [49]. All code is publicly available on GitHub https://github.com/NeuroCSUT/ViraMiner.

## 5 Acknowledgements

Supported by the Nordic Information for Action eScience Center awarded by the Nordic Acadey for Advanced Sciences (NordForsk), the Swedish Research Council and the Swedish Cancer Society.

